# Tactile perception of auditory roughness

**DOI:** 10.1101/2022.10.16.512429

**Authors:** Corentin Bernard, Richard Kronland-Martinet, Madeline Fery, Sølvi Ystad, Etienne Thoret

**Affiliations:** Aix-Marseille University, CNRS, UMR7061 PRISM, Marseille, 13009, France

**Keywords:** Roughness, Beating, Audio-tactile

## Abstract

Auditory roughness resulting of fast temporal beatings is often studied by summing two pure tones with close frequencies. Interestingly, the tactile counterpart of auditory roughness can be provided through touch with vibrotactile actuators. However, whether auditory roughness could also be perceived through touch, and whether they exhibit similar characteristics is unclear. Here, auditory roughness perception and its tactile counterpart were evaluated using similar two pure tones stimuli. Results revealed similar roughness curves in both modalities suggesting similar sensory processing. This study attests of the relevance of such a paradigm for investing auditory and tactile roughness in a multisensory fashion.

## 1. Introduction

Auditory roughness is a fundamental acoustical attribute due to very fast fluctuations of sounds that conveys emotions (Arnal et al., 2015), drives musical consonance (Helmholtz, 1885; Plomb & Levelt, 1965), and shapes orchestral timbres (Pressnitzer & McAdams, 2000). It is also at the basis of the definition of critical bands (Terhardt, 1974), a fundamental property of cochlear filters characterizing the ability of the cochlea to separate two pure tones. Auditory roughness has therefore led to a significant body of work in psychoacoustics to identify the acoustic factors that modulate this auditory sensation.

To unveil the mechanisms underlying auditory roughness perception, studies have used combinations of monochromatic tones (Miśkiewicz et al., 2006), pure tones, leading to models of perceived auditory roughness (Daniel et al., 1997; Vassilakis, 2001; Leman, 2000). A combination of two monochromatic tones can indeed create amplitude modulations that induce a sensation of roughness depending on the frequency spacing between the two tones, also defined by the frequency ratio between the two frequencies. The roughness of such sounds increases until reaching a maximum and then decreases as the frequency ratio between the two frequencies increases, as presented in Fig. 1. Since auditory roughness is often described as characterizing very fast fluctuations in sounds, such stimuli are perfect candidates to investigate such a sensation.

**Fig. 1.**
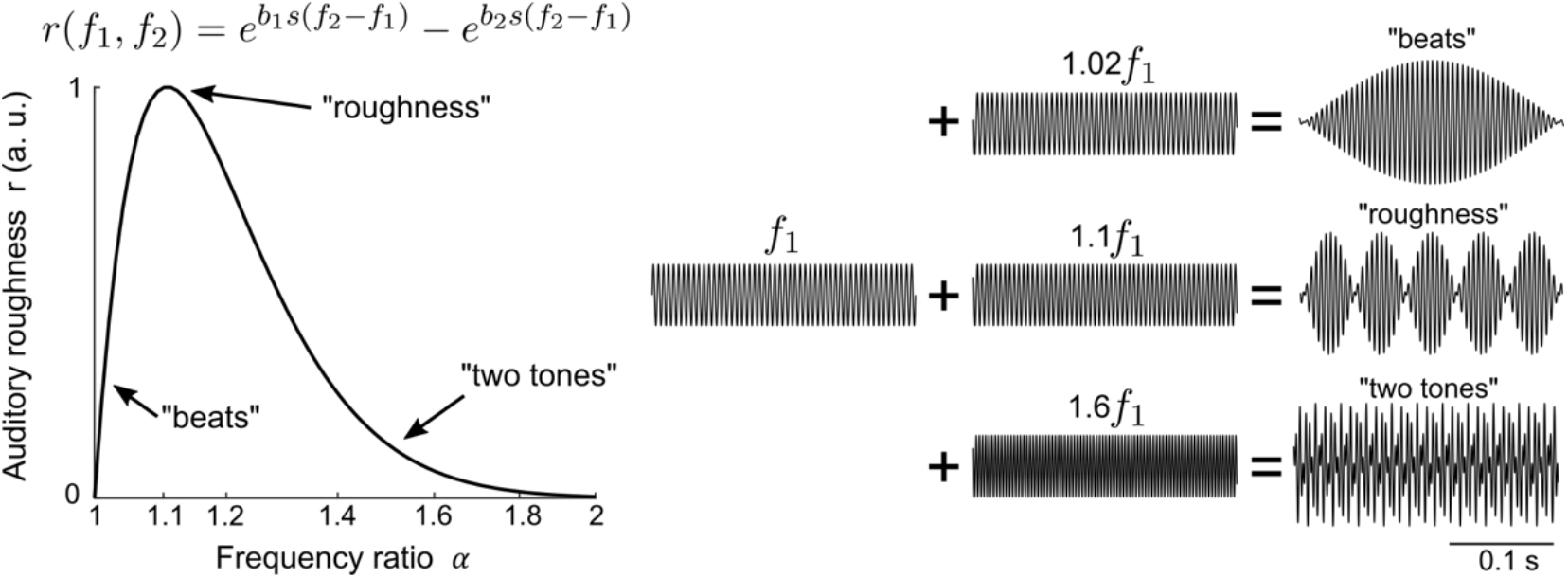
Typical auditory roughness perception according to the frequency ratio of a sum of two pure tones: 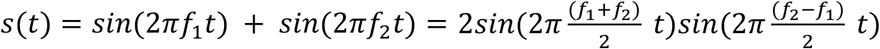. When the frequency ratio *α*=*f*_2_/*f*_1_ is small, the combination of tones tends to be perceived as one tone slowly modulated by the other one, a sensation that is characterized as beats. When the frequency ratio increases, a sensation of roughness appears. As *α* becomes even larger, the perceived roughness decreases and the two tones are perceived separately. This theoretical auditory roughness curve is defined with the parameters *b*_1_ = 3.5, *b*_2_ = 5.75, 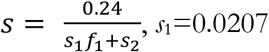 and *s*_2_ = 18.96 (Vassilakis, 2001).

On the other hand, for the tactile modality, surface roughness refers to one of the principal perceptual attributes used to describe textures explored with the finger (Tiest & Kappers, 2006). Perceived surface roughness is defined by the physical and geometrical properties of the textures, such as the height and the density of the asperities (Taylor & Lederman, 1975; Stevens & Harris, 1962). Typical stimuli for surface roughness evaluation are sandpapers with various grit sizes. Interestingly, similar observations showing that the perceived surface roughness depends on the spatial frequency and the amplitude of the haptic signal were made on synthetic textures with new haptic touchscreen technologies (İşleyen et al., 2019; Bodas et al., 2019).

In a recent review on multimodal roughness perception, Di Stefano & Spence (2022) made a clear distinction between auditory roughness and surface roughness: auditory roughness is a temporally based perceptual property, that is experienced through hearing, while surface roughness is spatiotemporal one, related to textures, that is assessed by touch (and vision). However, whether the sensation of auditory roughness as historically investigated could also be experienced through touch is still unclear.

In this paper, we investigate the tactile counterpart of auditory roughness using the same two pure tones signals in both modalities. In the auditory domain, stimuli were presented through headphones while a vibrotactile actuator was used in the tactile domain. The experiment was carried out in two sessions, one with auditory stimuli and the other with tactile stimuli. The goal was to determine the relationship between the roughness sensation and the frequency ratio of two different pure tones to compare the roughness curves in both modalities. Showing such a coherence would suggest that hearing and tactile sensory systems share similar processes. This would also validate these stimuli for further investigations in a multisensory context.

## 2. Methods

### 2.1 Participants

18 participants (8 women, M=30 years old. between 21 and 57 years old), 15 right-handed, voluntarily took part in the experiment. None of them reported having any auditory problems or skin concerns. The participants gave their informed consent before the experiment. The experiment lasted about 1 hour. The experimental protocol was validated by the local ethical committee.

### 2.2 Stimuli

The experiment was composed of two sessions: one auditory and one tactile. The same signals were used for both modalities and the stimuli were constructed by combining two monochromatic tones: *s*(*t*) = *sin*(2*πf*_1_*t*) + *sin*(2*πf*_2_*t*) of frequencies *f*_1_ and *f*_2_=*αf*_1_. Four frequency conditions were considered (*f*_1_ = 50, 100, 200, 300 Hz). The experimental design did not include higher frequency conditions since frequencies above 800 Hz are not perceptible by the human tactile sensory system (Verrillo, 1969). The frequency ratio *α* ranged from 1 to 2. When *α*=1, *f*_1_=*f*_2_ the tones are at unison, when *α*=2, *f*_1_=2*f*_2_, the tones are separated by one octave. Twelve values of *α* were chosen (1, 1.01, 1.02, 1.03, 1.05, 1.10, 1.15, 1.20, 1.25, 1.35, 1.50, 2.00). For each frequency condition, stimuli with different *α* values were compared pairwise leading to 66 pairs. Overall, each subject performed 2 sessions with 4 blocks of 66 pairs which led to 528 trials.

### 2.3 Apparatus

Sounds were presented through Sennheiser HD-650 headphones at a sampling rate of 44100 Hz powered by a Pioneer A-209R audio amplifier. Tactile stimuli were presented through an Actronika HapCoil-One vibrotactile actuator (dimensions: 11.5 × 12 × 37.7 mm3, acceleration: 8 g-pp, Frequency bandwidth: 10 to 1000 Hz, resonant frequency: 65 Hz). This kind of actuator has already been used in the literature to render the sensation of textures with vibrations (Rocchesso et al., 2016). The actuator was powered by a Pioneer A-209R audio amplifier. The subjects were asked to grab the vibrotactile actuator between the thumb and the index of their right hand. During the tactile experiment, participants wore noise-canceling headphones to prevent them from using potential auditory cues produced by the tactile device to perform the task.

### 2.4 Task and procedure

Participants were seated in front of a computer screen in a quiet room. They started randomly either with the audio or tactile session. The experiment was a pairwise comparison. At each trial, two stimuli with different values of *α* were presented successively. The stimuli lasted for 1 second and pairs were sequentially presented with an inter-stimulus interval of 800 ms. For each pair of stimuli, the participants were asked to determine which stimulus was the most “granular” (granuleux in French). We avoided the terms “rough” and “pleasant” that are commonly used in the literature since several participants had a musical background which might have influenced their judgment. For each subject, the 4 blocks and the stimuli within each block were presented in a randomized order and for each pair, the presentation order was also randomized. Responses were collected with a keyboard and the interface was designed with Max/MSP software to provide either audio or tactile stimuli. The volume and the intensity of auditory and tactile stimuli were set constant during the whole experiment.

### 2.5 Data analysis

Based on the subject’s responses, each stimulus was assigned a score of perceived auditory roughness. This score is the ratio of the number of times subjects judged the stimulus as rougher by the number of times the stimulus was presented in the pairwise comparison (=11). For one subject, if a stimulus of a given *α* value has been judged N times as more “granular” than another, its roughness score equals N/11. Finally for each subject, each modality and each *f*_1_ condition, roughness curves characterizing the roughness score evolution with respect to the *α* value were computed.

## 3. Results

Firstly, the results showed that the roughness curves obtained were coherent with the theoretical roughness curve proposed by Vassilakis (2001) and Leman (2000). This validates the present protocol for roughness score measurement (correlations between the 8 mean curves, 2 modalities x 4 frequency ratio condition, and the theoretical curves: r(9) : M=0.89, range=[min=0.83, max=0.97], all p<0.001), see Fig. 2.

**Fig. 2.**
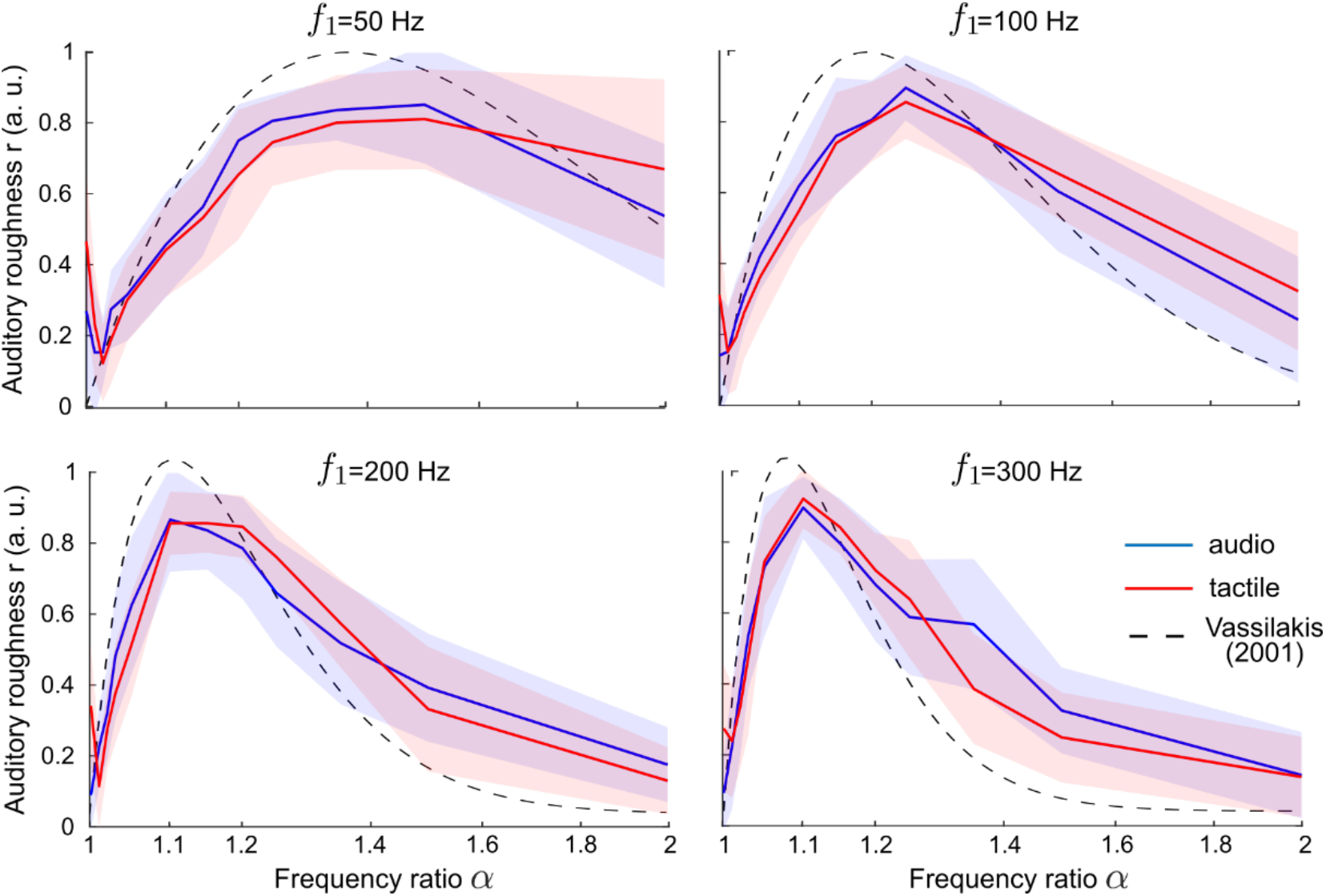
Audio (blue) and tactile (red) roughness curves obtained from the experiment for the four frequency conditions. Mean across subjects is presented in solid lines, and standard deviation is presented with the shaded zones. The black dashed lines depict the theoretical auditory roughness model proposed by Vassilakis (2001).

Most importantly, the results revealed that auditory and tactile modalities provide similar roughness curves in the 4 frequency conditions tested, see Fig. 2. We observed a significant correlation between the tactile and auditory roughness mean curves for the 4 conditions (Pearson correlations: *f*_1_=50 Hz : r(9)=0.94, p=4.7.10^−6^, BF10=5e^−3^; *f*_1_=100 Hz: r(9)=0.97, p=2e^−7^, BF10=9e^4^; *f*_1_=200 Hz: r(9)=0.92, p=1.7e^−5^, BF10=1.7e^3^; *f*_1_=300 Hz: r(9)=0.94, p=3.9e^−7^, BF10=6.5e^3^).

Lastly, in order to assess potential differences between the two modalities, we computed the differences between auditory and tactile roughness scores for each frequency condition *f*_1_ and *α* values and for each subject. This resulted in a population of 864 samples, whose mean does not significantly differ from 0 (t-test: t_863_=1e^−15^ p=1, CI95%=[-0.0149; 0.0149]), showing that there is no significative difference between the audio and tactile roughness curves.

## 4. Discussion

In this paper, auditory roughness perception induced by temporal beatings was compared between the auditory and the tactile sensory systems. In one experiment with two sessions, subjects had to compare the roughness of pairs of sounds or tactile vibrations parametrized by the ratio between the frequencies of two monochromatic components. Roughness curves were then computed for both modalities and were found to be similar in the two modalities and with theoretical auditory roughness model.

Auditory roughness curves are coherent with the model of Vassilakis (2001). Interestingly, we observe the same curves in the tactile modality. In particular, the position of maximal roughness perception changes according to the lower frequency *f*_1_. It would therefore be of interest now to further investigate the multimodal process of auditory roughness perception. It may indeed lead to a common process between the two modalities and the critical band framework (Makous et al., 1995) could also be relevant in the tactile domain. In addition, a formal comparison with the modulation transfer function in the amplitude modulation domain could also be done with specific experiments (Weisenberger, 1986).

Secondly, this study further sheds light on more fine similarities in the temporal processing of vibrations through these two modalities. One major difference between the two modalities is that for high *α* values (around 2), audition discriminates the two harmonic frequencies leading to a perception of two tones, whereas in touch, we perceive only one uniform complex temporal vibration (Bensmaia, & Hollins, 2000).

It is worth noticing that the auditory roughness curves present similar shapes to the surface roughness curves (Unger et al., 2010) by making analogies between sound beating frequency and texture spatial frequency (i.e., the number of ridges per millimeters). Firstly, for low spatial frequencies, only large undulations of the texture are felt. The surface roughness then increases as the spatial frequency increases until reaching a maximum. Lastly, surface roughness decreases as the ridge’s density highly increases until the texture becomes almost smooth. Although auditory and surface roughness remain two distinct perceptual attributes, there could hereby be processed with similar underlying mechanistic processes. In addition, recent evidence has shown that rhythm perception is shared between audio and haptics even for textures explored through active touch (Bernard et al., 2021). Our current findings suggest that these results could be extended to the perception of beating and temporal roughness.

## 5. Conclusion

This study provides results for the investigation of audio-tactile roughness under a common framework. The framework classically dedicated to investigating auditory roughness has here been formally validated for the tactile modality. One particularly relevant perspective is the investigation of the multisensory integration of these stimuli in congruent or incongruent situations. We may expect to observe audio-tactile interactions and how one modality can enhance the perception of roughness in the other. This has already been observed in several multisensory situations (Jousmäki & Hari, 1998, Guest et al., 2002, Yau et al., 2009). These cases are of great relevance to understanding the fine mechanistic bases of human perceptual systems.

## Acknowledgments

This work was supported by an ILCB/BLRI grant ANR-16-CONV-0002 (ILCB), ANR-11-LABX-0036 (BLRI), the Excellence Initiative of Aix-Marseille University (A*MIDEX), the Sound and Music from Interdisciplinary and Intersectorial Perspectives (SAMI - A*MIDEX) project and France Relance. The authors would like to thank MIRA, Aflokkat and Nicolas Huloux for his thoughtful comments on the paper.

